# Monolayer-Crystal Streptavidin Support Films Provide an Internal Standard of cryo-EM Image Quality

**DOI:** 10.1101/091355

**Authors:** Bong-Gyoon Han, Zoe Watson, Jamie H. D. Cate, Robert M. Glaeser

**Author notes:** Corresponding author: Robert M. Glaeser, Lawrence Berkeley National Laboratory, University of California, Berkeley, CA 94720.

## Abstract

Analysis of images of biotinylated *Escherichia coli* 70S ribosome particles, bound to streptavidin affinity grids, demonstrates that the image-quality of particles can be predicted by the image-quality of the monolayer crystalline support film. The quality of the Thon rings is also a good predictor of the image-quality of particles, but only when images of the streptavidin crystals extend to relatively high resolution. When the estimated resolution of streptavidin was 5 Å or worse, for example, the ribosomal density map obtained from 22,697 particles went to only 9.5 Å, while the resolution of the map reached 4.0 Å for the same number of particles, when the estimated resolution of streptavidin crystal was 4 Å or better. It thus is easy to tell which images in a data set ought to be retained for further work, based on the highest resolution seen for Bragg peaks in the computed Fourier transforms of the streptavidin component. The refined density map obtained from 57,826 particles obtained in this way extended to 3.6 Å, a marked improvement over the value of 3.9 Å obtained previously from a subset of 52,433 particles obtained from the same initial data set of 101,213 particles after 3-D classification. These results are consistent with the hypothesis that interaction with the air-water interface can damage particles when the sample becomes too thin. Streptavidin monolayer crystals appear to provide a good indication of when that is the case.

## INTRODUCTION

Affinity support films have a number of features that are attractive for preparing specimens for single-particle electron cryo-microscopy (cryo-EM for short). It is thought that affinity binding should be generally more structure-friendly than adsorption of biological macromolecules to continuous carbon films, and certainly less hazardous than interaction with the air-water interface within open holes. In addition, binding to an affinity substrate offers the possibility of achieving a high density of particles per unit area, even when the particle concentration is relatively low, such as ~40 nM (Han et al., 2016).

A number of different types of affinity grid have been introduced in recent years. One of these is based on Ni-NTA lipid monolayers that are picked up on holey-carbon films (Benjamin et al., 2016; Kelly et al., 2010; Kelly et al., 2008). Another is based on chemical functionalization that can be performed on oxidized, continuous carbon films (Llaguno et al., 2014). A more recent idea has been to simply bind antibodies non-specifically to a carbon support films, and then use these as affinity grids (Yu et al., 2016).

Using streptavidin (SA) monolayer crystals to make affinity grids represents yet another approach (Crucifix et al., 2004; Han et al., 2016; Wang et al., 2008). There are numerous ways to take advantage of the high binding affinity of biotin, and also of streptavidin-binding peptides, to immobilize particles of interest onto the SA support film. Examples include the use of biotinylated “adaptor molecules” (Crucifix et al., 2004); random chemical biotinylation of lysine residues on the surface of proteins (Han et al., 2012); and – for membrane proteins – incorporation into proteoliposomes that include a biotinylated lipid (Wang and Sigworth, 2010). A further feature of the SA affinity grid is that the image of the SA crystal can be subtracted by Fourier filtering, when desired (Wang et al., 2008).

We now report that a further benefit of using monolayer crystals of SA is that they provide a convenient way to assess single-particle cryo-EM image quality. The image-quality of the SA lattice itself is easily evaluated, because the signal is confined to reciprocal lattice points (“Bragg spots”). By comparison, it takes much more effort to evaluate the image quality of single particles because of the lengthy process involved in obtaining a high-resolution density map. It thus is quite useful that the image of the SA lattice provides a good indication of the quality of images of single particles bound to the lattice.

To illustrate the use of SA crystals as an indicator of single-particle image quality, we obtained 3-D reconstructions of *Escherichia coli* 70S ribosomes using images in which the resolution of the SA lattice was either (1) no better than 5 Å or (2) 4 Å or better. Importantly, the quality of the Thon rings was the same in both cases. When the images of SA crystals extended to only 5 Å or worse, the resolution of the density map obtained from 22,697 particles extended to only 9.5 Å. On the other hand, when the resolution of the SA lattice was better than 4 Å, the density map from the same number of particles extended to a resolution of 4.0 Å. The resolution of the map improved even further, to 3.6 Å, when 57,826 particles from such images were used.

## MATERIALS AND METHODS

The data set analyzed in this study is the same one that was collected as part of our initial description of how to make long shelf-life SA affinity grids (Han et al., 2016). To summarize, biotinylated *E. coli* 70S ribosomes, which had been incubated with 20 μM spectinomycin, were bound to carbon-backed SA monolayer crystals that spanned the open holes of Quantifoil grids. Cryo-EM images were collected at 300 keV with a Gatan K2 camera, using a low-base FEI Titan electron microscope, equipped with a Gatan 626 cold holder. LEGINON (Suloway et al., 2005) was used to automate the data collection. The movie frames were re-aligned and summed with the most recent version of software developed at UCSF (Li et al., 2013), referred to as MotionCor2. The ribosome particles used in the current work were the same ones that were boxed previously.

A key step added in the present work consisted of using an automated method to quantitatively characterize the quality of images of SA crystals. The first step in this process was to unbend the images of SA lattices, using scripts provided in the 2dx package (Gipson et al., 2007). Another script, written in MATLAB (The MathWorks, Inc.), was then used to define the resolution zone within which the average IQ value, as reported in 2dx after unbending, increased to a value of 7.

In addition, the values reported by CTFFIND4 (Rohou and Grigorieff, 2015), for the cross correlation between the experimental Thon rings and the corresponding theoretical models of the CTF, were used as a second measure of the quality of the image. In this instance, the cross correlation was limited to the region between 1/(8 Å) and 1/(4 Å), and Bragg spots from the SA lattice were first removed, using the script for Fourier filtering described in (Han et al., 2016).

Single-particle data-processing of images of ribosome particles was done with tools in the RELION(Scheres, 2012) package, version 1.4. Final refinements were always carried out with a mask placed around the large ribosomal subunit. Further details of the processing are mentioned in the Results section, below.

## RESULTS

Although all holes in the carbon film that were queued up for automated data collection appeared to be equally good when viewed in SEARCH mode, the corresponding high-magnification images of streptavidin were not all of equal quality. After unbending (Henderson et al., 1986) the streptavidin lattice, the highest resolution at which Bragg spots with IQ values of or 4 could be seen extended beyond 4 Å in many, but not all, of the images. For other images, however, Bragg spots with IQ values of 3 or 4 did not go beyond 10 Å. Examples of these two cases are shown in Figure 1. The initial interpretation might be that some of the streptavidin crystals simply were “not good to begin with”. However, a deeper analysis of the data, presented below, indicates that this interpretation is not correct.

**Figure 1.**
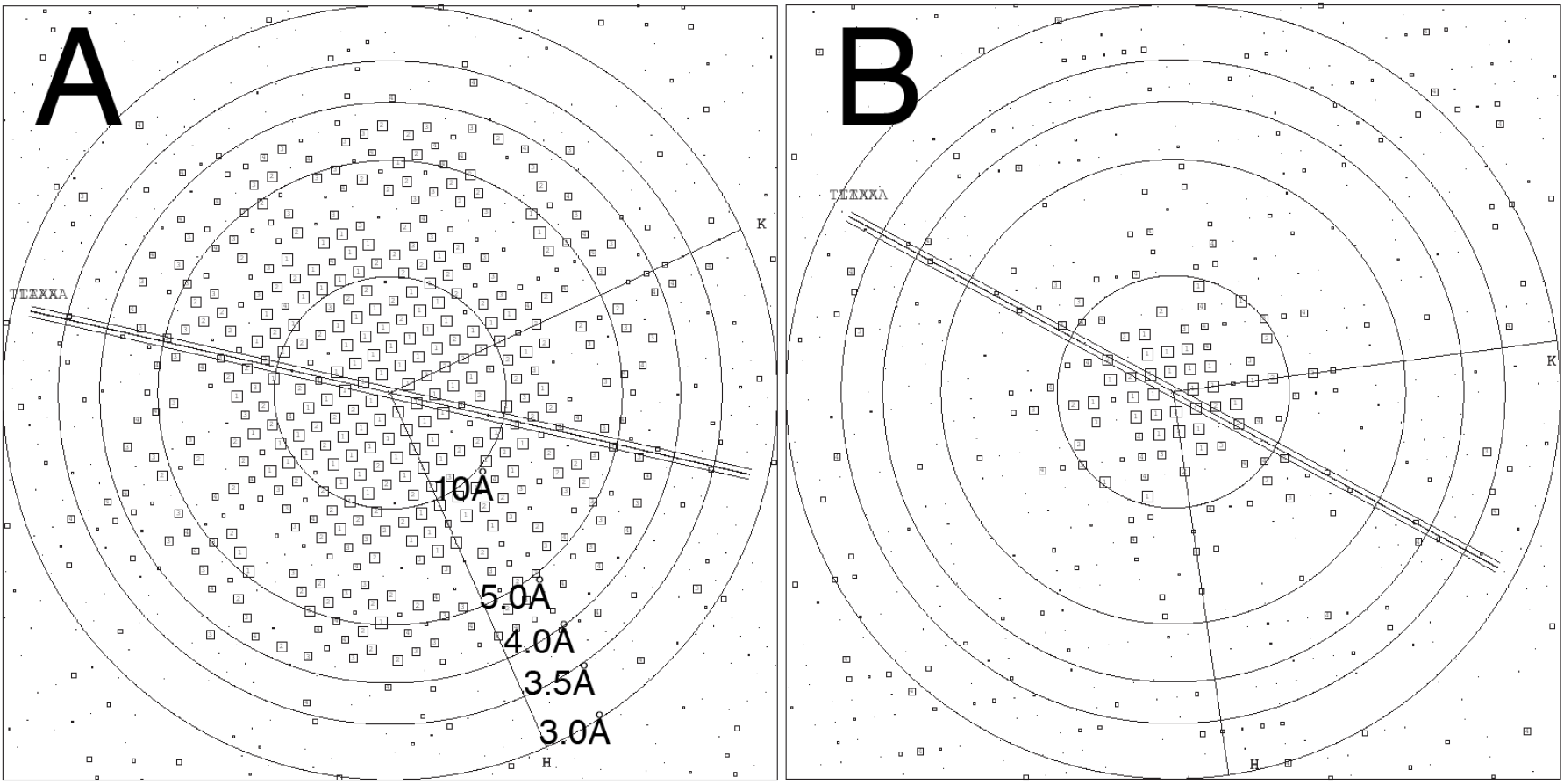
Representative examples of IQ plots produced after unbending images of streptavidin monolayer crystals. Circles of different radius are superimposed at different resolution values, as labeled in panel A. (A) The IQ plot for an image selected from within the middle of the distribution of “good images” that is shown in Figure 2B. Diffraction spots with IQ values of 4 or less, which are shown as larger boxes, extend to a resolution of 4 Å, with a few being present at even higher resolution. (B) The IQ plot for an image selected from within the middle of the distribution of “poor images” that is shown in Figure 2B. Diffraction spots with IQ values of 4 or less extend only to a resolution somewhere between 5 Å and 10 Å.

In order to estimate objectively the image quality of each SA crystal, we first plotted the reciprocal of the values of IQ, averaged within successive resolution shells, as a function of resolution. The IQ value of individual Bragg spots in unbent images was first introduced as a figure of merit to evaluate the quality of the data by (Henderson et al., 1986). When IQ = 7, the estimated phase error is 45° - see Table 10.1 in (Glaeser et al., 2007), and the contributions of signal and noise within such a Bragg spot are equal.

Figure 2A shows examples of such plots, using the two images whose Fourier transforms are shown in Figure 1. Also shown in Figure 2A is the fit of a Gaussian function to the data that is of the form

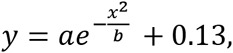

where *y* is the reciprocal of the average value of IQ in a given resolution zone, and *x* is the resolution corresponding to the midpoint of each such zone. The fixed constant, 0.13, is an estimate of the reciprocal of the average IQ, in the output provided by 2dx, when there is little or no signal in the Bragg spots.

**Figure 2.**
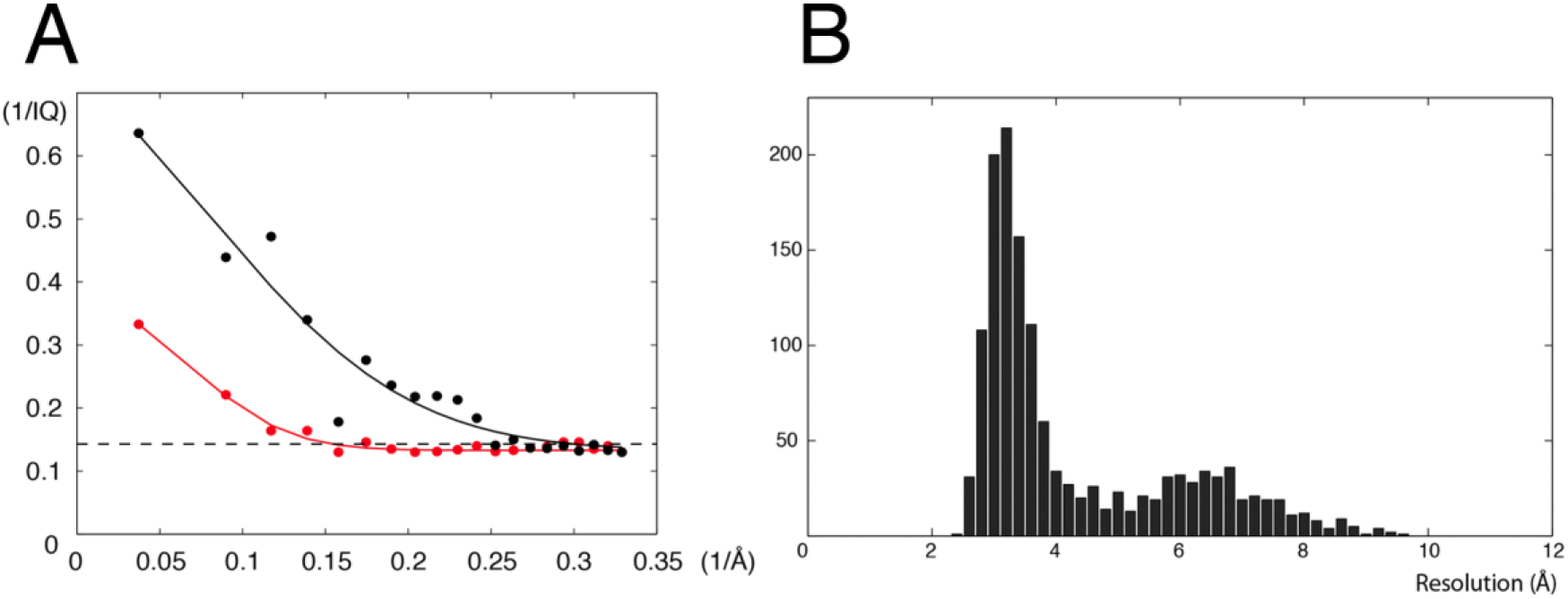
Explanation of how the IQ plot is used to evaluate the quality (i.e. resolution) of an image. (A) The values of IQ after unbending, as reported by 2dx, are first averaged within successive resolution shells. The reciprocal of this average value is then plotted as a function of spatial frequency. Finally, a Gaussian function, defined in Equation 1, is fitted to the data. The resolution of the image of the SA crystal is then estimated to be the value at which the fitted curve falls to 1/7. (B) Histogram showing the number of images in resolution bins versus the resolution. The bimodal distribution seen here was not anticipated.

The resolution at which 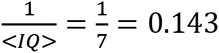 for the fitted curve is then used as an indication of the quality of the image of the streptavidin crystal. Unexpectedly, the distribution of the quality (i.e. “resolution”) of streptavidin images was found to be bimodal, as is shown in Figure 2B. There is a relatively narrow peak in the distribution centered at an estimated resolution of ~3.5 Å, and a second, relatively broad peak centered at a resolution of ~6.5 Å.

We first wanted to understand why the distribution of images with respect to their quality was bimodal. The simplest possibility would be that “some grids are good and some are not”, due to unknown factors in the many steps in their preparation. The situation turns out to be more complicated. As is illustrated in Figure 3, the quality of images of the streptavidin crystals can vary within one grid, as well as from one grid to another. Furthermore, when there is variation within one grid, there seem to be local areas that are good and others that are bad, as is suggested by there being sequential “runs” with similar image quality.

**Figure 3.**
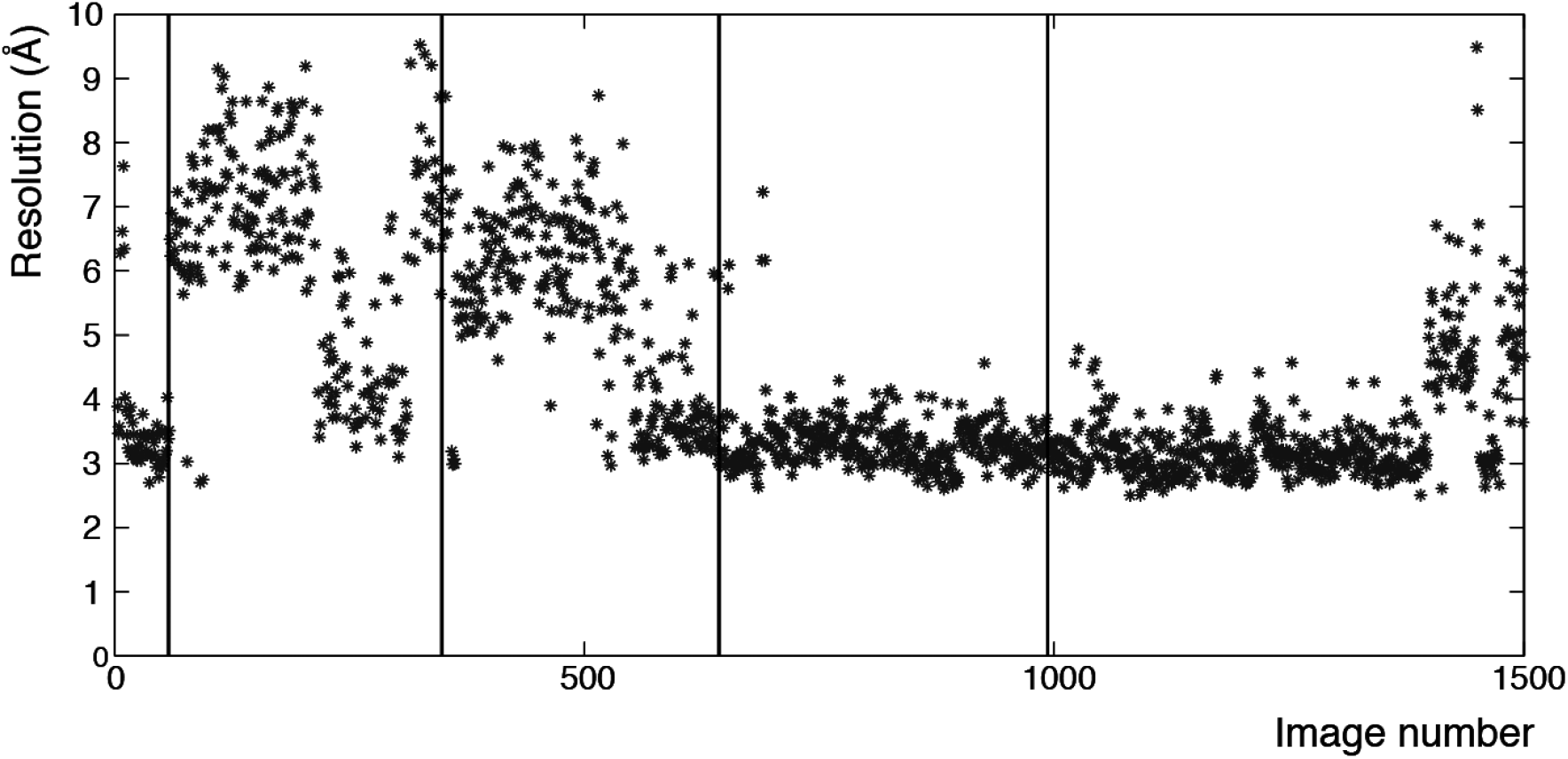
Plot of the estimated resolution of the SA crystal in a given image as a function of the sequential number given to that image within the full data set. Vertical lines indicate boundaries between images that were recorded from different grids.

In addition, the expectation that the resolution of the SA crystals would scale with the resolution of the Thon rings is also incorrect. Instead, as is shown in Figure 4, a 2-D scatter plot of SA-image quality vs Thon-ring quality again exhibits a bimodal distribution. It is informative that the images lying within the banana-shaped data-cloud do, in fact, go to somewhat higher resolution when the Thon rings are also better, and, conversely, the resolution values for the SA images within this group are somewhat worse when the Thon rings are not as good. On the other hand, images within the second cloud of data points, for which the resolution of the SA lattice is poor, mainly exhibit quite good Thon rings.

**Figure 4.**
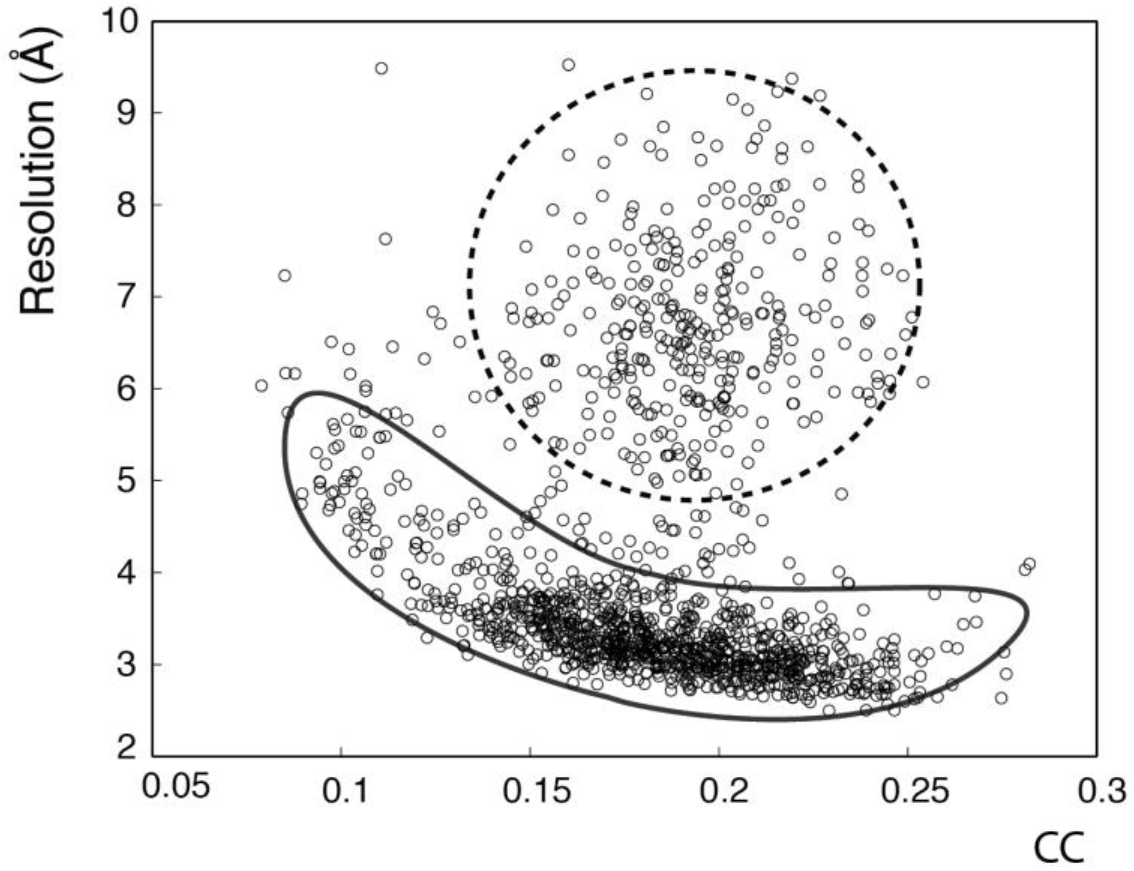
Scatter plot of the estimated resolution of the SA crystal versus the cross-correlation between the continuous Fourier transform of the image and a model of the CTF for the defocus value of that image. CTF fitting, and the corresponding values of the cross-correlation value in the resolution zone between 8 Å and 4 Å, were obtained by using CTFFIND4 (Rohou and Grigorieff, 2015). The plot shows a bimodal distribution, with two clouds indicated schematically by dotted and solid outlines for “bad” and “good” SA crystals, respectively.

We next investigated whether the data-quality for ribosome particles would correlate with the data-quality for the SA crystals to which they were bound. To do this, we created two subsets of particles, depending upon where their images fell with respect to the two clouds in the scatter plot shown in Figure 4. More specifically, one group consisted of 22,697 particles for which the resolution for the SA crystals was 5 Å or worse, and the second group consisted of the same number of particles for which the resolution for the SA crystals was 4 Å or better. We then obtained refined density maps for the two groups. As is shown in Figure 5, the resolution of the map was only about 9.5 Å for particles in the first group, i.e. ones boxed from images in which the resolution of the SA lattice was 5 Å or worse. On the other hand, the resolution obtained for the second group was quite good (4.0 Å), considering the relatively small number of particles that was used to obtain the density map.

**Figure 5.**
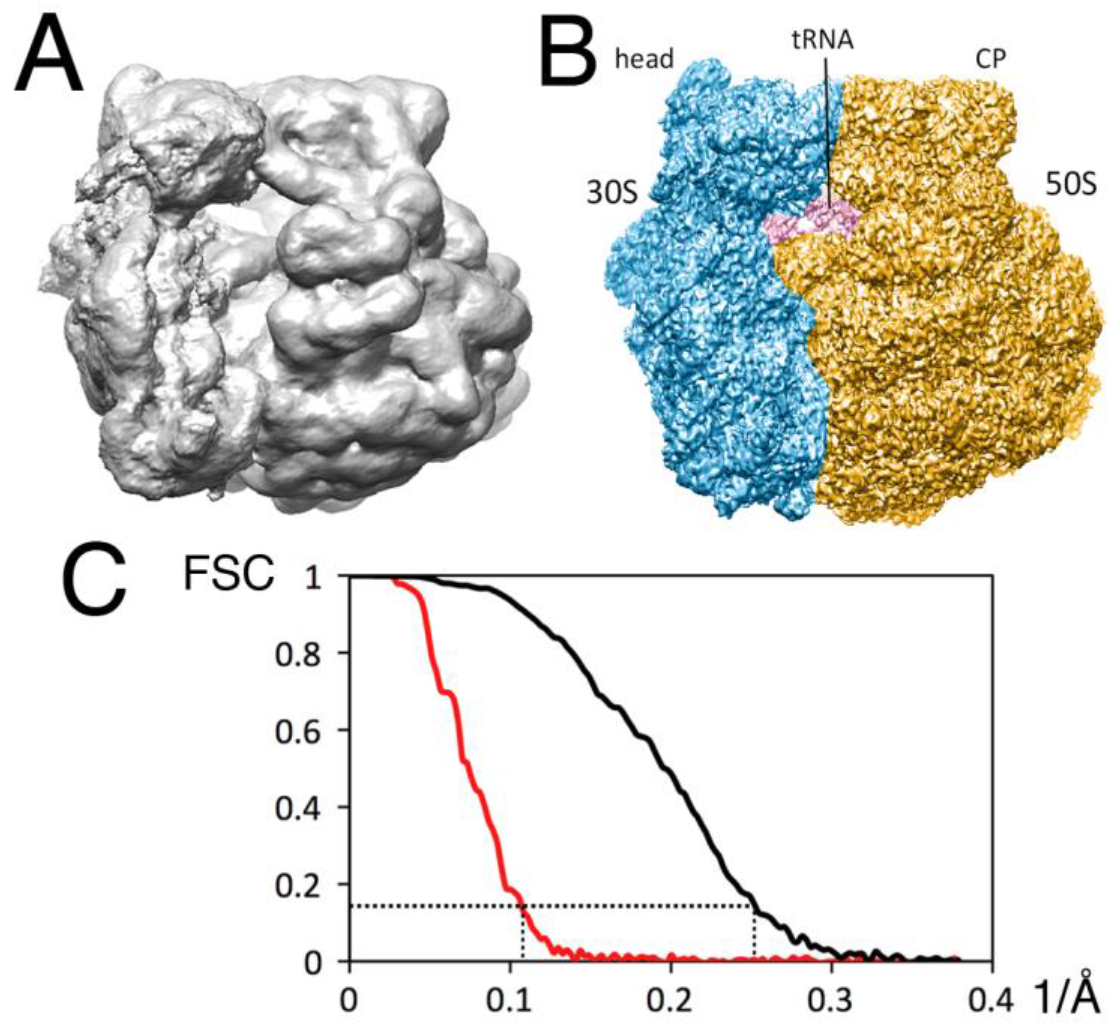
Result obtained when performing refinement using the same number of particles boxed from good images and from bad images, respectively. (A) The density map for 22,697 ribosomes boxed from bad images. (B) The density map for the same number (22,697) of ribosomes boxed from good images. Head, head domain of the 30S subunit; CP, central protuberance of the 50S subunit. (C) The “gold standard” FSC curves for the two maps shown in panels (A) and (B) respectively; the red curve is for ribosomes boxed from bad images and the black curve is for ribosomes boxed from good images. The point where the value of the FSC curve is 0.143 is indicated by the dashed lines.

Next, we determined the resolution obtained when using the full set of ribosome particles boxed from “good” images. To do this, we first edited that data set by performing 3 rounds of 2- D classification, deleting, in each round, mainly those particles for which the class averages showed no significant evidence that they contained 70S ribosomes. This process reduced the number of particles from 61,646 to 57,826. We then used RELION to calculate a refined (single-class) density map. As is shown in Figure 6, the resolution of the map, determined by the “gold standard” FSC, was 3.6 Å. This resolution was the same, to two significant figures, for the refined map obtained from the unedited data set of 61,646 particles, confirming that the candidate particles removed by editing contained little value of interest.

**Figure 6.**
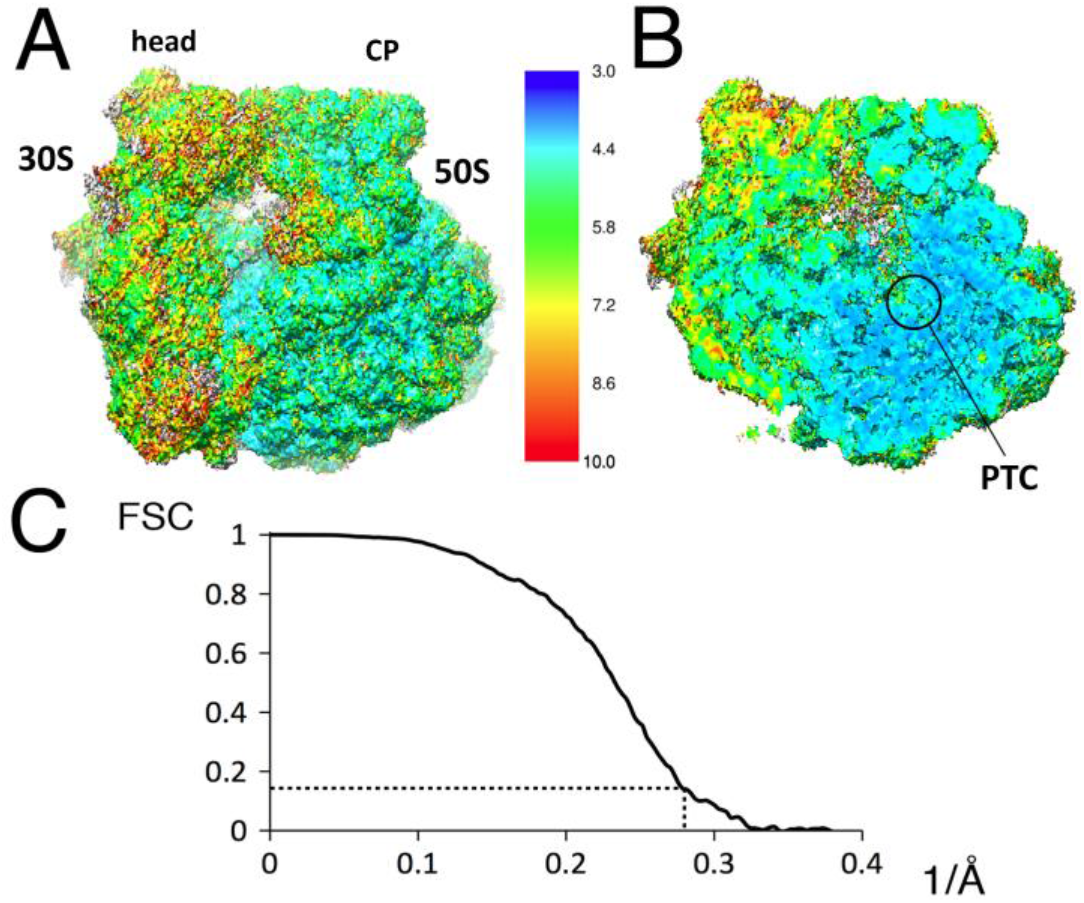
Map obtained when using ribosomes picked from good SA images. The particles were treated as being in a single class, i.e. 3-D classification was not used to address the expected conformational heterogeneity. As stated in Methods, a mask surrounding the 50S subunit was used during refinement. (A) Refined map for particles in the edited set of good images. Head, head domain of the 30S subunit; CP, central protuberance of the 50S subunit. (B) A cutaway of the local-resolution representation of the density map, for a plane that includes the peptidyl transferase center (PTC) of the 50S subunit. (C) The corresponding “gold-standard” FSC curve for the map shown in (A) and (B). The point where the value of the FSC curve is 0.143 is indicated by the dashed lines.

Encouraged by the relatively high resolution obtained, we performed 3-D classification of particles in the edited data set. We classified these particles into only three classes because the size of the data set was relatively small. As is shown in Figure 7, this produced two major classes consisting of 70S ribosome particles and a third, minor class consisting mainly of 50S particles. Refinement of the two major classes resulted in structures at resolutions of 4.3 Å (from 14,848 particles) and 3.8 Å (from 35,937 particles), respectively.

**Figure 7.**
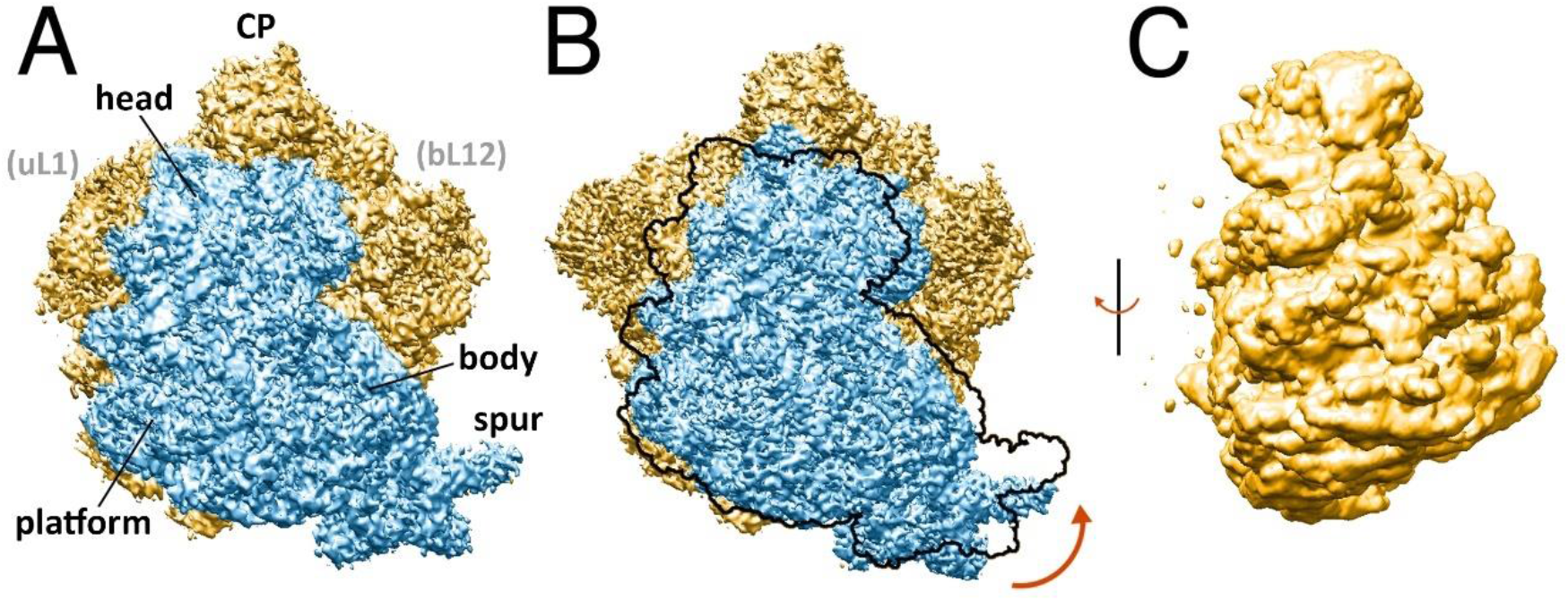
Results obtained after using 3-D classification to sort particles, which were all obtained from “good” images, into three smaller but more homogeneous subsets. (A) Density map for 14,846 particles assigned to one of the classes, representing an unrotated state of the ribosome. Features of the ribosome are indicated, including the location of uL 1 and bL 12 of the 50S subunit, which are not visible in the present map due to disorder. (B) Density map for 35,937 particles assigned to a second class. The outline of the subvolume of the 30S subunit, in the position occupied in panel A, is superimposed on the density map shown in B. The well-known rotation of the 30S subunit (Frank and Agrawal, 2000) is inticated by the curved arrow. (C) Density map for 7,041 particles in the third class shows that it consists mainly of 50S particles. The resolution after 3D refinement was 4.3 Å, 3.8 Å and 8.2Å in panels (A), (B), and (C), respectively.

Comparison of the density maps produced by particles in the two major classes shows the familiar intersubunit “rotation” (Frank and Agrawal, 2000) of the 30S particle relative to the 50S particle. This figure was prepared by first using Chimera (Pettersen et al., 2004) to align the maps in panels (A) and (B) such that their 50S densities overlapped well with one another. Each 70S map was then compared to the X-ray crystal structures available as PDB accession numbers 4V9D and 4YBB. Map A was found to correlate best with the rotated P/E state in 4V9D assembly I, while map B correlates best with the unrotated state in 4YBB assembly 2.

## DISCUSSION

Our analysis of the quality of cryo-EM images of ribosome particles prepared on SA-affinity grids is based on three quantitative metrics. One metric is the estimated resolution limit for the image of the SA-crystal. A second metric is based on the Thon rings seen in the Fourier transforms of images. Parenthetically, we believe that the thin carbon film that was evaporated onto the lipid-tail side of the SA crystals should make the main contribution to the Thon rings. The third metric is the resolution of the three-dimensional reconstruction obtained for biotinylated 70S ribosomes that were bound to the affinity grids.

Discarding poor-quality images reduces the computational burden involved during refinement while, at the same time, it is expected to improve the final result. As the data in Figure 4 demonstrate, however, it is not sufficient to use just the Thon rings alone to select images with the best quality. Instead, significant value is also added by discarding images in which the diffraction peaks from the SA crystals fail to go to high resolution. The criterion of SA image quality thus complements other reasons for discarding images such as “high drift, low signal, heavy ice contamination, or very thin ice”, mentioned by (Loveland et al., 2016).

The bimodal distribution that we found for the resolution of SA crystals – see Figure 2 and Figure 4 – indicates that something other than excessive amounts of inelastic scattering can be a strong factor affecting image quality. The broad peak in Figure 2, centered at a resolution of ~6.5 Å, obviously comprises mainly those images that form a diffuse, circular cloud in Figure 4. Importantly, as already mentioned, these same images tend to be quite good when one looks at the Thon rings, indicating that the ice was relatively thin for all of them.

Turning attention first to the banana-shaped cloud in Figure 4, the resolution values of the SA images does become slightly worse as the quality of Thon rings become worse, as might be expected. We hypothesize that the simultaneous loss of resolution for SA crystals and for the Thon rings may well be due, in this case, to increasing ice thickness. It is possible to test this hypothesis in the future by recording images with a microscope that is equipped with a zero-loss energy filter, because the number of electron counts in the image would be measurably less, the thicker the specimen.

Focusing attention next on the bimodal distribution seen in Figure 4, it is significant that the ribosome particles appear to be damaged when the images of the SA crystals are poor, as is shown by the lower resolution of the density map that is obtained from such particles (compare Figure 5A to Figure 5B). One possible explanation is that the sample in such images – both ribosomes and streptavidin – was either partially dehydrated or in some other way contacted by the air-water interface after blotting. It is puzzling, however, that images in which the sample is good and those in which it is bad can both occur over a wide range of specimen thicknesses, as inferred from the quality of the Thon rings. A possible explanation is that the sample was spared from being damaged by a surfactant monolayer at the air-water interface in the good images, whereas in “bad images” the surfactant – if any – was too dilute to be effective. Since this hypothesis is complicated and speculative, however, it will not be easy to test it.

An alternative explanation notes that the background contributed by the SA layer must be less perfectly subtracted when the crystals are disordered. The question that then arises is whether this extra amount of background might cause alignment and assignment of Euler angles to be less accurate, thereby causing the refined density map for 22,697 particles to extend to only 9.5 Å. In order to test this alternative, we boxed 22,697 areas of background from images in which the SA was poor, and added these background-boxes, one at a time, to ribosome particles boxed previously from images in which the SA component went to a resolution of 4 Å or better. These intentionally corrupted images of ribosome particles were then used to obtain a refined density map, which was found to extend to a resolution of 4.8 Å. This value of resolution is only slightly worse than the value of 4.0 Å obtained from the same, limited number of particles before corrupting the images with additional noise. This result seems to rule that out the hypothesis that added background, resulting from imperfect subtraction of the SA component, is responsible for the significantly lower resolution obtained from images in which the Thon rings are excellent but the SA component is not.

Finally, it is still to be determined whether the carbon-backed version of SA affinity grids can be used for high-resolution imaging of macromolecular particles that are significantly smaller than ribosome particles. While the SA “background” can be subtracted from images by Fourier filtering, the noisy signal due to the carbon film is likely to eventually interfere with boxing particles and with alignment and assignment of Euler angles as the particle size decreases. The same limitation applies to making cryo-specimens directly on continuous carbon films, of course. We thus point out that the option may exist to use single-layer graphene to “back” SA monolayer crystals. An attractive approach to do so, not yet tested, would be to grow SA crystals on the surface of small wells, as described in (Wang and Sigworth, 2010), and pick these up with grids already covered by graphene. While picking up monolayer crystals with evaporated carbon films does not perform very well (Kubalek et al., 1991), graphene may be more hydrophobic than evaporated carbon, and thus may perform better. What we have tried, in preliminary experiments, is to pick up lipid monolayers on graphene coated grids, and then use the on-grid method to grow SA crystals. The lipid transfer appeared to work well, as did the binding of SA to biotinylated lipid. Nevertheless, only small crystalline domains were obtained, possibly because the fluidity of lipids adsorbed to graphene may be much less than that of lipids spanning open holes in holey carbon films.

## ACKNOWLEDGMENTS

This work was supported in part by NIH grants P01 GM051487 and R01 GM065050, and by the National Science Foundation Graduate Research Fellowship Program under Grant number 1106400. Any opinions, findings and conclusions or recommendations expressed in this material are those of the authors and do not necessarily reflect the views of the National Science Foundation. Molecular graphics and analyses were performed with the UCSF Chimera package. Chimera is developed by the Resource for Biocomputing, Visualization and Informatics at the University of California, San Francisco (supported by NIH grant P41 GM103311).

